# CD1b-specific T cells are transcriptionally closer to conventional CD4 T cells than to innate-like NKT and MAIT cells

**DOI:** 10.64898/2026.08.25.747105

**Authors:** Alan Hsieh, Kattya Lopez, Segundo R León, Roger I Calderon, Leonid Lecca, Megan B. Murray, D. Branch Moody, Sara Suliman, Ildiko Van Rhijn

## Abstract

Unconventional T cells recognize non-peptide antigens presented by molecules other than the major histocompatibility complex (MHC) proteins. Among unconventional T cells, natural killer T (NKT) cells, which recognize CD1d-lipid complexes, mucosal-associated invariant T (MAIT) cells, which recognize MR1-metabolite complexes, and γδ T cells, are thoroughly studied. CD1b presents self- and mycobacterial lipids to relatively understudied T cell subsets. Like MAIT cells and type I NKT cells, CD1b-specific cells include subpopulations with conserved TCRs. Consequent to their recognition of a nearly monomorphic antigen-presenting molecule, CD1b-specific T cells might share innate-like features with MAIT, type I NKT, and γδ T cells. Due to their low frequency in the peripheral blood, CD1a-, CD1b-, and CD1c-specific T cells have been studied predominantly as in vitro-expanded clones, so even basic information about their native ex vivo immunophenotypes is lacking. Here, we sort and transcriptionally profile ex vivo two T cell populations that recognize CD1b presenting mycobacterial mycolipids and compare them with conventional CD4 and CD8 T cells, γδ T cells, NK cells, MAIT cells, and NKT cells. We show that both the invariant TCR-expressing CD4^+^, CD1b-GMM-specific germline encoded mycolyl-reactive (GEM) T cells, as well as the diverse TCR-expressing CD1b-GMM-specific T cells, are transcriptionally closer to conventional T cells than to the innate-like T cell populations γδ, MAIT and type I NKT cells. Thus, despite their recognition of non-polymorphic antigen presenting molecules, CD1b-specific T cells show adaptive rather than innate-like transcriptional features.

## Introduction

Conventional CD4⁺ and CD8⁺ T cells express diverse T cell receptors (TCRs) and are activated by recognition of pathogen-derived peptide antigens presented by polymorphic antigen presenting molecules. For normal function, clonal expansion and differentiation into effector and memory populations is needed prior to mediating systemic response, pathogen clearance and long-term protective immunity. Because the priming and activation of naïve conventional T cells requires at least a week to reach full effect (1) early responses by the innate immune system are important. Although T cells are traditionally regarded as mediators of adaptive immunity, several subsets display innate-like characteristics. These include mucosal-associated invariant T (MAIT) cells, natural killer T (NKT) cells, and Vδ2Vγ9 gammadelta (γδ) T cells, which circulate in the body with an activated- or effector phenotype characterized by rapid effector function upon TCR triggering, responsiveness to inflammatory cytokines independent of T cell receptor (TCR) engagement, and expression of the transcription factor promyelocytic leukemia zinc finger (PLZF), which drives the innate-like T-cell transcriptional program (2, 3).

In addition to the innate-like phenotype, MAIT cells, NKT cells, and Vδ2Vγ9 gammadelta (γδ) share the fact that they do not recognize peptide-MHC through cognate interaction, so they also fit the definition of unconventional T cells. Instead, they interact with other proteins on antigen presenting cells, including, in many cases, non-polymorphic MHC-like molecules, which also led to the designation donor-unrestricted T cells (DURTs) (4, 5). Since the antigen-binding clefts of non-polymorphic MHC-like molecules are specialized to accommodate non-peptide ligands, the T cells that recognize them respond to non-peptide antigens, including lipids and metabolites. Previous work established an “innateness” gradient that organizes adaptive, innate-like, and innate lymphocytes according to their transcriptional profiles, placing conventional CD4 T cells at the adaptive extreme and NK cells at the innate extreme. This continuum captures the gradient from transcriptional programs supporting clonal expansion and proliferative capacity of T cells normally present at low precursor frequency versus those that exist at higher precursor frequency that prioritize immediate effector function and rapid immune responsiveness (6).

In humans, the non-polymorphic MHC-like antigen-presenting molecules are MR1, CD1a, CD1b, CD1c, and CD1d. Among the CD1a, CD1b, CD1c, and CD1d-specific T cells, many are autoreactive, where they either recognize CD1 molecules themselves regardless of the bound antigen, or they recognize CD1 molecules bound to common self-lipids (7–12). There are also CD1a, CD1b, CD1c, and CD1d-specific T cells that only recognize CD1 molecules bound to exogenous, rare (glyco)lipids (13–16). Examples of this category include germline-encoded mycolyl-reactive (GEM) T cells, and LDN5-like T cells, which are both highly specific for CD1b presented mycolipids, a class of lipids only found in a subset of mycobacteria, including *Mycobacterium tuberculosis* and the tuberculosis (TB) vaccine Bacille-Calmette Guérin (BCG) (17, 18).

Because CD1d and MR1 present antigens to NKT- and MAIT innate-like T cell populations, respectively, T cells specific for CD1b, which is also non-polymorphic, are generally assumed to also be innate-like. However, whether CD1b-specific T cells function primarily as innate-like responders to inflammatory cues or whether they are predisposed to mediate durable antigen-specific immunological memory has not been systematically tested. This knowledge gap has arisen because CD1b reactive T cells have mainly been studied as long term clones and lines rather than ex vivo T cell populations (18–23).

Especially for GEM- and LDN5-like T cells this is a relevant question because they recognize mycobacterial lipids and might therefore contribute to *M. tuberculosis* clearance during TB. On the one hand, GEM- and LDN5-like T cells resemble conventional T cells in that they are present at low precursor frequencies in peripheral blood and undergo antigen-driven proliferation in response to pathogen-derived antigens encountered during infection, supporting their ability to mediate memory responses (24, 25). On the other hand, supporting their grouping with NKT and MAIT cells, GEM T cell express T cell receptor (TCR) motifs that, although using a different Variable (V)-Joining (J) segment combination and a different CDR3 motif, show the typical signs of an invariant TCR α chain and a biased use of a TCR β chain variable (V) segment (17). Additional CD1b-mycobacterial lipid-specific T cells (LDN5-like cells) show V segment bias but no clear CDR3 motif and lastly, some do not have any of these TCR biases and have a more heterogeneous TCR repertoire (8). Here, we ask whether CD1b-mycobacterial lipid-specific T cells exhibit a canonical innateness program or resemble adaptive CD4 and CD8 T cells leveraging samples from a cohort of mycobacteria-exposed participants.

## Materials and methods

### Tetramers and antibodies

Long chain synthetic methoxy MA (26) and short chain GMM isolated from *Rhodococcus equi* were used to load biotinylated CD1b monomer (NIH tetramer facility) according to previously published methods (24, 27, 28). PE-labeled MA- and PE-labeled GMM-loaded CD1b tetramers were mixed 1:1 before staining. Ready-to-use CD1d-PBS-57 tetramer-BV421and MR1-5-OP-RU tetramer-APC were from the NIH tetramer facility.

### Cell lines

As positive controls for CD1b-GMM tetramers, primary LDN5 T cells (29) were spiked into PBMC. As positive control for the CD1b-MA tetramers, Jurkat cells transduced with the TCR of MA-specific T cell clone 28 (28) were used (Jurkat.TCRclone28).

### Human participants

Members of a previously reported tuberculosis cohort (24) were re-recruited for another blood donation based on previously detected high levels of CD1b tetramer^+^ T cells in their PBMC. Demographic data of the subjects included in this study are provided in **Table 1**. Participants with a positive QuantiFERON-TB Gold In-Tube (QFT) assay and no clinical evidence of active tuberculosis were classified as having latent *Mycobacterium tuberculosis* infection (QFT^+^), whereas participants with a negative QFT result were classified as uninfected (QFT^−^). We also included four participants with culture-confirmed pulmonary TB at the time of first inclusion, who were being treated or had finished treatment at the time of re-inclusion. For validation of reagents, PBMC isolated from de-identified leukoreduction filters from local blood bank donors, provided by the Brigham and Women’s Hospital Specimen Bank, were used. The Institutional Review Board of the Harvard Faculty of Medicine and Partners Healthcare, and the Institutional Committee of Ethics in Research of the Peruvian Institutes of Health approved this study protocol. All adult study participants were adults and provided informed consent.

**Table 1:** Demographic data and cell numbers per sequenced population. Demographic data of participants and respective numbers of cells that were sorted for low input RNA sequencing are shown. Subjects in italics were used for studies of reproducibility and the effect of population size on outcome. The cell numbers in grey font are populations of which the RNAseq data did not pass quality control.

|  | demographic data |  |  | numbers of cells in each RNA-profiled population |  |  |  |  |  |  |  |  |
| --- | --- | --- | --- | --- | --- | --- | --- | --- | --- | --- | --- | --- |
| participant | gender | age | TB status | CD4 | CD8 | NK | MAIT | NKT | Vd1 | Vd2 | GEM | non GEM<br>CD1b specific |
| 157-5 | female | 40 | QFT<sup>+</sup> | 1000 | 1000 | 1000 | 810 | 1000 | 1000 | 850 | 93 | 769 |
| 158-5 | female | 37 | QFT<sup>+</sup> | 1000 | 1000 | 1000 | 0 | 0 | 1000 | 0 | 0 | 905 |
| 159-8 | male | 28 | QFT<sup>+</sup> | 1000 | 1000 | 0 | 1000 | 1000 | 1000 | 1000 | 0 | 0 |
| 160-9 | female | 43 | QFT<sup>-</sup> | 1000 | 1000 | 1000 | 1000 | 1000 | 1000 | 1000 | 31 | 626 |
| 208-9 | male | 43 | QFT<sup>+</sup> | 1000 | 1000 | 0 | 1000 | 0 | 0 | 1000 | 152 | 558 |
| 209-8 | female | 36 | QFT<sup>+</sup> | 1000 | 1000 | 1000 | 1000 | 1000 | 0 | 0 | 0 | 0 |
| 195-4 | male | 20 | post-TB | 1000 | 1000 | 1000 | 1000 | 1000 | 1000 | 651 | 0 | 0 |
| 196-3 | female | 22 | post-TB | 527 | 1000 | 1000 | 1000 | 1000 | 1000 | 1000 | 0 | 0 |
| 083-0 | male | 60 | post-TB | 1000 | 1000 | 1000 | 1000 | 1000 | 1000 | 1000 | 0 | 0 |
| 197-0 | male | 66 | post-TB | 1000 | 1000 | 1000 | 1000 | 1000 | 1000 | 1000 | 429 | 0 |
| *085-9* | *male* | *28* | *unknown* | *50, 50, 1000* | *0* | *50, 50, 1000* | *0* | *0* | *0* | *50, 50, 1000* | *0* | *0* |
| *110-6* | *female* | *44* | *unknown* | *50, 50, 1000* | *0* | *50, 50, 1000* | *0* | *0* | *0* | *50, 50, 1000* | *0* | *0* |
| *118-8* | *female* | *64* | *unknown* | *50, 50, 1000* | *0* | *50, 50, 1000* | *0* | *0* | *0* | *50, 50, 1000* | *0* | *0* |

### Cell sorting and handling

PBMC were isolated from 50 ml of blood and cryopreserved. Before sorting, cells were thawed, washed and resuspended in PBS with 2% BSA, all four tetramers were added for 10 min at room temperature; unlabeled anti-CD3 antibody clone OKT3 was added for 5 min at room temperature, antibody mix was added for 30 min at 4C. Cells were washed and resuspended in PBS with 2% BSA. The antibody mix consisted of: CD3-BV786; CD8-BV510; (both from BD biosciences); CD4-AF700; CD14-APC-Cy7; CD19-APC-Cy7; CD56-BV711; TCRαβ-PE-Cy7; Vα7.2-BV605 (specific for TRAV1-2); Vδ2-FITC (all from Biolegend), in Brilliant Stain buffer (BD biosciences). Cells were sorted on a 5-laser FACSAria II flow cytometer (BD Biosciences) and the collected data was analyzed with FlowJo software.

### RNA-seq analysis

RNA-seq library preparation was performed at the Broad Technology Labs (Broad Institute) using a modified Smart-seq2 protocol for low-input RNA-sequencing, which improves detection of full-length transcripts using template switching and pre-amplification. Samples from different participants and cell subsets, as well as water blanks, were randomized on the 96-well plates. Sequencing was performed in an Illumina NextSeq500 as paired-end 2 x 25 bp reads with an additional 8 cycles for each index.

We processed RNA-seq data on the UCSF Wynton high-performance computational cluster, using STAR alignment (30), trimmomatic to remove poor quality sequencing reads (31), then Kallisto for gene quantification to generate gene count matrices. Count data was analyzed using the DESeq2 package in R Studio (version 2026.01.1) (32).

## Results

### Study enrollment

We previously quantified mycolic acid (MA)- and glucose monomycolate (GMM)-loaded CD1b tetramer-specific T cells and MAIT cells in a cohort of BCG-vaccinated participants with TB disease and their household contacts with and without latent *Mtb* infection in Lima, Peru (24, 33). Because we did not observe differences in the peripheral blood frequencies of these subsets across TB disease states, we re-recruited a subset of participants with detectable frequencies of CD1b-specific T cells, irrespective of TB status (n=13, **Table 1**). The median age of participants was 40 years (interquartile range [IQR], 28–44 years), and 54% of participants were female. Among the re-recruited participants were four participants whom we had previously studied during active TB disease, when we profiled CD1b-specific T cells in the peripheral blood (24). If indicated, participants had completed anti-TB treatment in accordance with the Peruvian national guidelines before enrollment in the present study. Hence, all participants were exposed to mycobacterial lipid antigens through vaccination with BCG and known exposure to *Mtb*. For each of the participants in the study, the demographic data, cell populations, and numbers of cells obtained after the second sort are listed in **Table 1**.

### Tetramer generation and validation

CD1b tetramers are not available as an antigen-loaded, tetramerized, and validated reagents, and some variability in in-house loading efficiency and background staining is expected (28). To validate the tetramers we spiked an established clonal CD1b-GMM-specific LDN5 T cell line (29) into peripheral blood mononuclear cells (PBMCs) from a random blood bank donor as a positive control for the GMM-loaded CD1b tetramer (**Figure 1A**). As a positive control for the MA-loaded CD1b tetramer, we stained Jurkat cells transduced with a previously described MA-specific TCR, clone 28 (**Figure 1B**) (28). The tetramers successfully stained their respective positive control cell types among non-relevant CD3-positive cells and were therefore suitable to specifically identify populations of CD1b-mycolipid-specific T cells.

**Figure 1:**
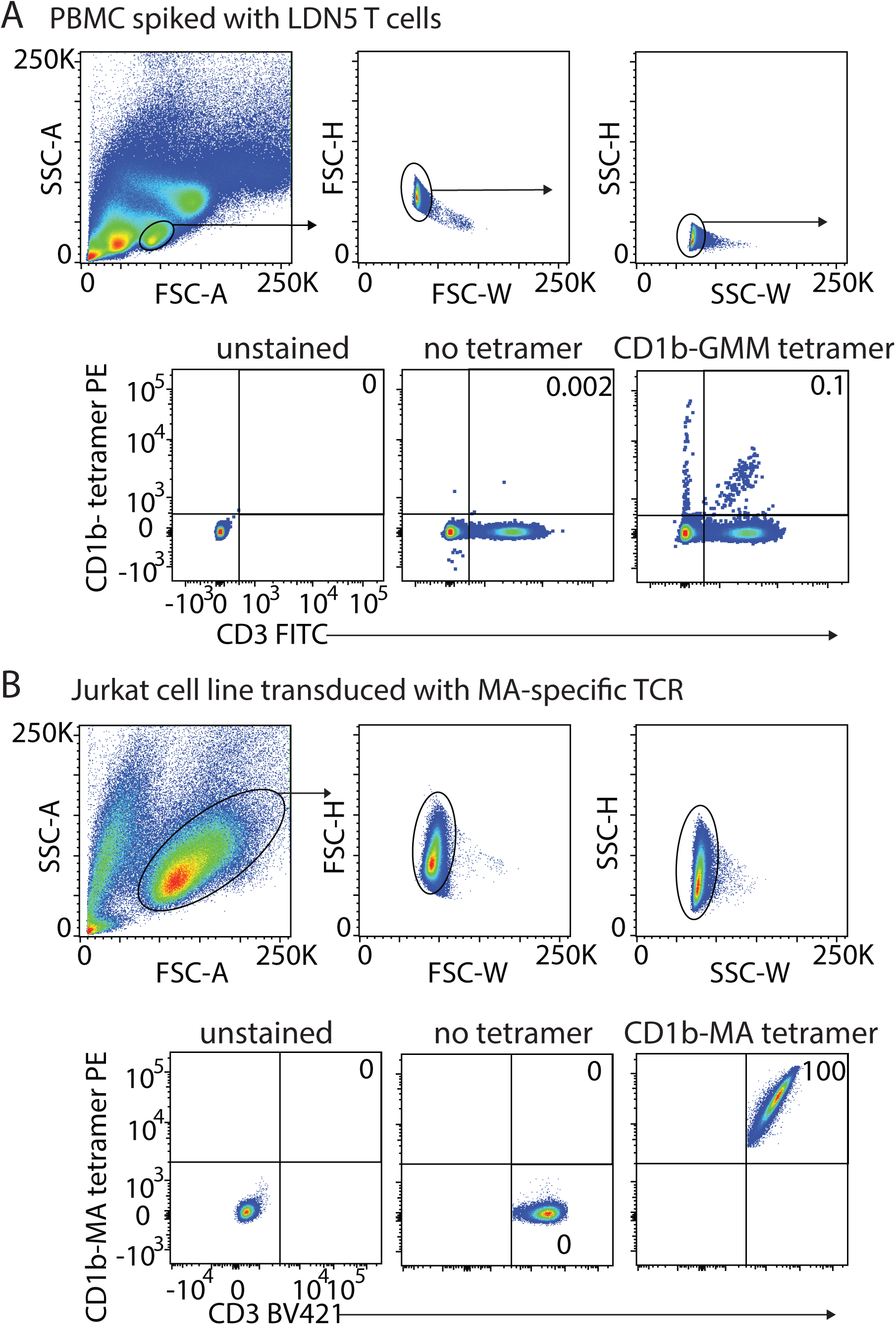
Validation of tetramers. **A** CD1b-GMM tetramers were validated using PBMC spiked with primary GMM-specific T cell line LDN5 T cells. **B** CD1b-MA tetramers were validated using a Jurkat cell line transduced with the MA-specific clone 28 TCR as a positive control.

### Cell populations and sorting

We then proceeded to define the transcriptional programs of CD1b-mycolipid-specific GEM T cells and other CD1b-mycolipid-specific T cells (called non-GEM CD1b-specific throughout the paper) in PBMCs. Due to the low numbers of these cells and the limited availability of PBMCs, we decided to perform low-input RNA sequencing (RNA-seq) of bulk-sorted CD1b tetramer^+^ cells (34). We used a mixture of CD1b-GMM and CD1b-MA tetramers for sorting because GEM T cells can recognize both antigens, and we wanted to maximize the numbers of sortable cells. For comparison with other T cell types, we also sorted conventional CD4 T cells, conventional CD8 T cells, NK cells, type I NKT cells, MAIT cells, Vδ2^+^ γδ T cells, Vδ2^−^ γδ T cells, capping the number of sorted cells at 1000 cells per population. We designed a gating scheme that, after gating for single live lymphocytes and excluding monocytes and B cells, in order to first separate CD3^−^ NK cells from T cells. From the CD3^+^ T cells, we gated MAIT cells using 5- (2-oxopropylideneamino)-6-d-ribitylaminouracil (5-OP-RU)-loaded MR1 tetramers, and type I NKT cells with CD1d tetramers loaded with the alpha galactosylceramide analogue (PBS-57). We then gated Vδ2^+^ and Vδ2^−^ γδ T cells from the remaining T cells. Vδ2^−^ γδ T cells in the blood mostly consist of Vδ1^+^ γδ T cells but can also include minor frequencies of Vδ3^+^ and other γδ T cells (35). Then, we gated CD1b tetramer^+^ cells from the remaining αβ T cells, which we separated into GEM T cells (TRAV1-2^+^, CD4^+^, GMM-loaded CD1b-tet^+^) and non-GEM CD1b-specific T cells (the remaining CD1b tetramer^+^ T cells), which included LDN5-like T cells and all other CD1b-mycolipid-specific T cells without defined TCR motifs. From all the tetramer^−^ T cells, we sorted conventional CD4 and CD8 single-positive cells (**Figure 2 and Supplementary Figures S1-9**). All sorted populations were then immediately re-sorted again based on the same markers to ensure the purity of rare subsets for transcriptional profiling (**Figure 2B**).

**Figure 2:**
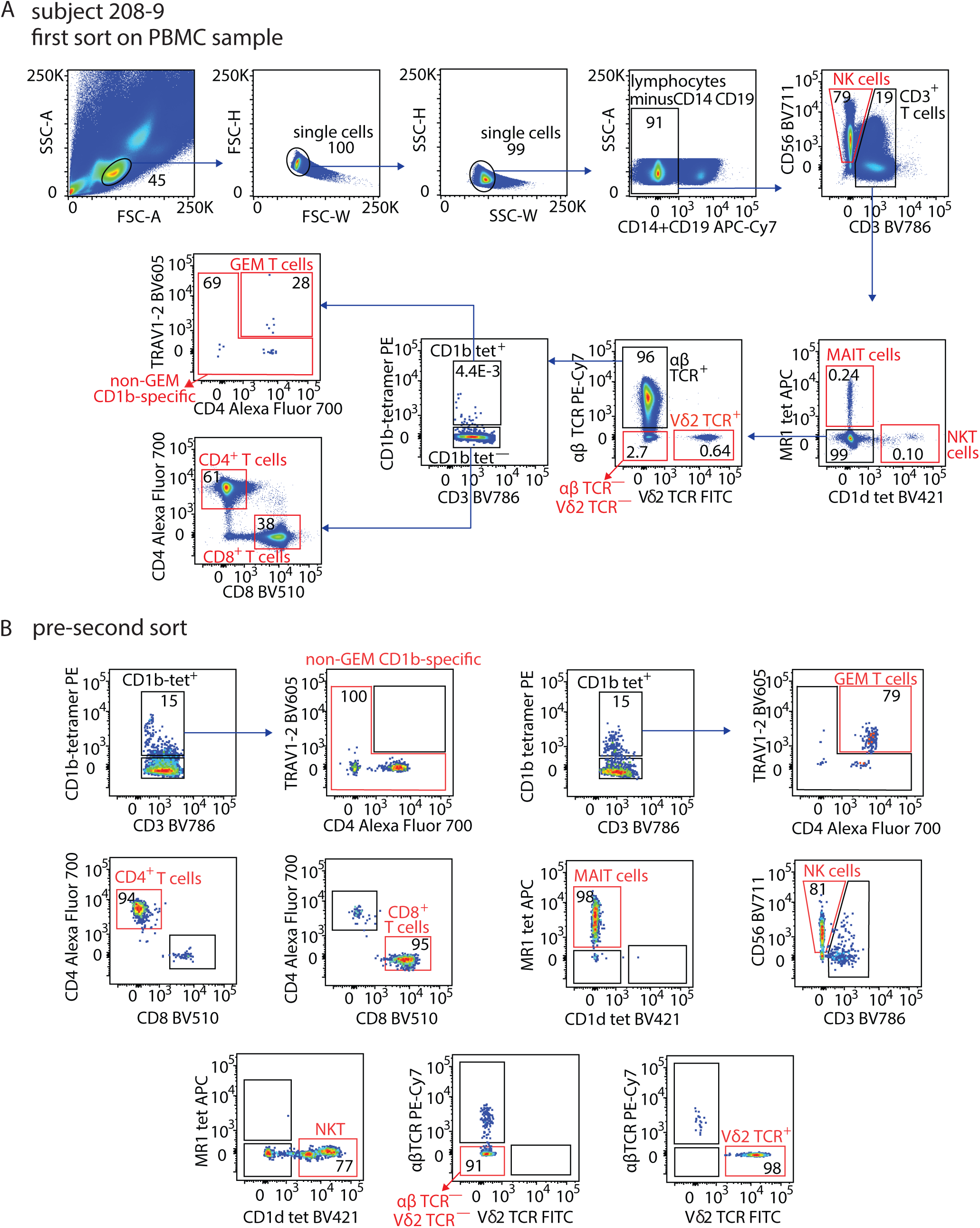
Gating scheme. **A** For the initial sort PBMC were first gated for single lymphocytes and removal of CD14+ monocytes and CD19+ B cells. NK cells (CD3^−^, CD56 dim or high) were separated from T cells (CD3+). From the T cells we gated MAIT (CD1d MR1-5-OP-RU tetramer^+^) cells and type I NKT cells (CD1d-PBS-57 tetramer^+^), and from the remaining T cells we gated αβTCR^−^ Vδ2^+^ γδ T cells and αβTCR^−^ Vδ2^−^ γδ T cells (consisting mostly of Vδ1^+^ γδ T cells). Then we gated the CD1b tetramer^+^ cells from remaining αβ T cells, which were then separated in GEM T cells (TRAV1-2^+^, CD4^+^, tetramer^+^) and non-GEM CD1b-specific T cells (the remaining tetramer^+^ cells). From the tetramer^−^ cells we sorted conventional CD4 and CD8 single postive cells. The red gates indicate the sorted populations. **B** Immediately after the first sort, a second sort was performed to obtain high purity cell populations. The red gates indicate the sorted populations.

### Quantification of T cell populations by flow cytometry

To assess the frequencies of the populations of interest as they occur in the PBMCs of our study participants ex vivo, before sorting, we analyzed the flow cytometry data that we collected during cell sorting (**Table 2**). Comparison with a published study in a cohort of healthy Australians on NK cells, Vδ2+ and Vδ2^−^ γδ T cells, NKT-, MAIT-, and GEM T cells (36) suggested that the population size we detected in the participants in our study fell within the expected range in a healthy population.

**Table 2:** Immunophenotyping of PBMC. Percentages of populations based on gating scheme in Figure 2 are shown. Subjects in italics were used for technical validation and studies of reproducibility and the effect of population size on outcome.

|  | in single lymphocyte gate |  |  | in CD3 gate |  |  |  |  |  |  |  |  |
| --- | --- | --- | --- | --- | --- | --- | --- | --- | --- | --- | --- | --- |
| subject | CD3 | NK | CD14+CD19 | CD4 | CD8 | MAIT | NKT | Vd1 | Vd2 | non-GEM<br>CD1b spec | GEM | all CD1b spec |
| **157-5** | 47 | 33 | 19 | 60 | 25 | 0.34 | 0.27 | 13 | 0.70 | 9.2E-03 | 2.9E-03 | 1.2E-02 |
| **158-5** | 70 | 5.2 | 22 | 55 | 24 | 0.17 | 0.17 | 17 | 1.3 | 2.5E-03 | 9.8E-05 | 2.6E-03 |
| **159-8** | 56 | 10 | 32 | 64 | 28 | 1.42 | 0.52 | 1.5 | 2.1 | 2.9E-03 | 4.7E-05 | 3.0E-03 |
| **160-9** | 56 | 14 | 28 | 74 | 20 | 0.18 | 0.066 | 3.5 | 0.77 | 1.0E-01 | 1.1E-02 | 1.2E-01 |
| **208-9** | 17 | 71 | 10 | 58 | 37 | 0.24 | 0.10 | 2.7 | 0.64 | 2.9E-03 | 1.2E-03 | 4.2E-03 |
| **209-8** | 68 | 9.1 | 21 | 59 | 34 | 0.56 | 0.25 | 2.6 | 1.5 | 3.8E-03 | 0 | 3.8E-03 |
| **195-4** | 56 | 23 | 17 | 63 | 30 | 0.70 | 0.12 | 2.2 | 2.2 | 2.3E-03 | 5.1E-05 | 2.4E-03 |
| **196-3** | 50 | 25 | 24 | 70 | 26 | 0.30 | 0.47 | 1.5 | 0.43 | 1.3E-03 | 0 | 1.3E-03 |
| **083-0** | 45 | 24 | 30 | 70 | 21 | 0.69 | 0.98 | 4.0 | 1.1 | 5.1E-03 | 7.2E-04 | 5.8E-03 |
| **197-0** | 31 | 41 | 27 | 53 | 37 | 0.30 | 0.10 | 2.1 | 5.5 | 6.6E-03 | 3.2E-04 | 6.9E-03 |
| ***085-9*** | *69* | *10* | *19* | *66* | *25* |  |  | *7.1* | *0.84* |  |  |  |
| ***110-6*** | *42* | *35* | *18* | *63* | *31* |  |  | *2.1* | *1.6* |  |  |  |
| ***118-8*** | *52* | *5.2* | *39* | *67* | *28* |  |  | *1.4* | *0.81* |  |  |  |

### Reproducibility and effect of population size on gene expression profile

While we were able to sort 1000 cells of most of the targeted cell populations, for some populations, notably the GEM and non-GEM CD1b-specific T cell populations, we were unable to obtain 1000 cells because of limited availability of PBMCs (**Table 1)**. To study technical reproducibility and the effect of population size in our RNA-seq experiments, we included three participants from whom we performed duplicate sorts of different numbers from otherwise identical populations (50 vs 1000 cells). To determine if there was an effect of the number of cells sorted on the overall transcriptional programs, we compared populations of 50 versus 1000 cells of CD4, Vδ2, and NK cells, and technical replicates of 50 cells of these populations from the three participants (**Table 1**, blue background). Although comparing samples generated from 50 versus 1,000 sorted cells identified differentially expressed genes, transcriptomes clustered primarily by cell subset and participant rather than by cell number. CD4, Vδ2, and NK cell samples from the same participant grouped regardless of the number of sorted cells, and technical replicates generated from 50 or 1,000 cells clustered closely. Thus, cell number had a lower impact on global transcriptional profiles relative to biological variation between cell subsets and participants. (**Figure 3A**). Therefore, although we recovered fewer CD1b-specific GEM and other T cells than their innate and adaptive counterparts (**Table 2)**, we do not expect the lower number of sorted cells to alter transcriptional profiles sufficiently to obscure the biological differences between cell subsets.

**Figure 3:**
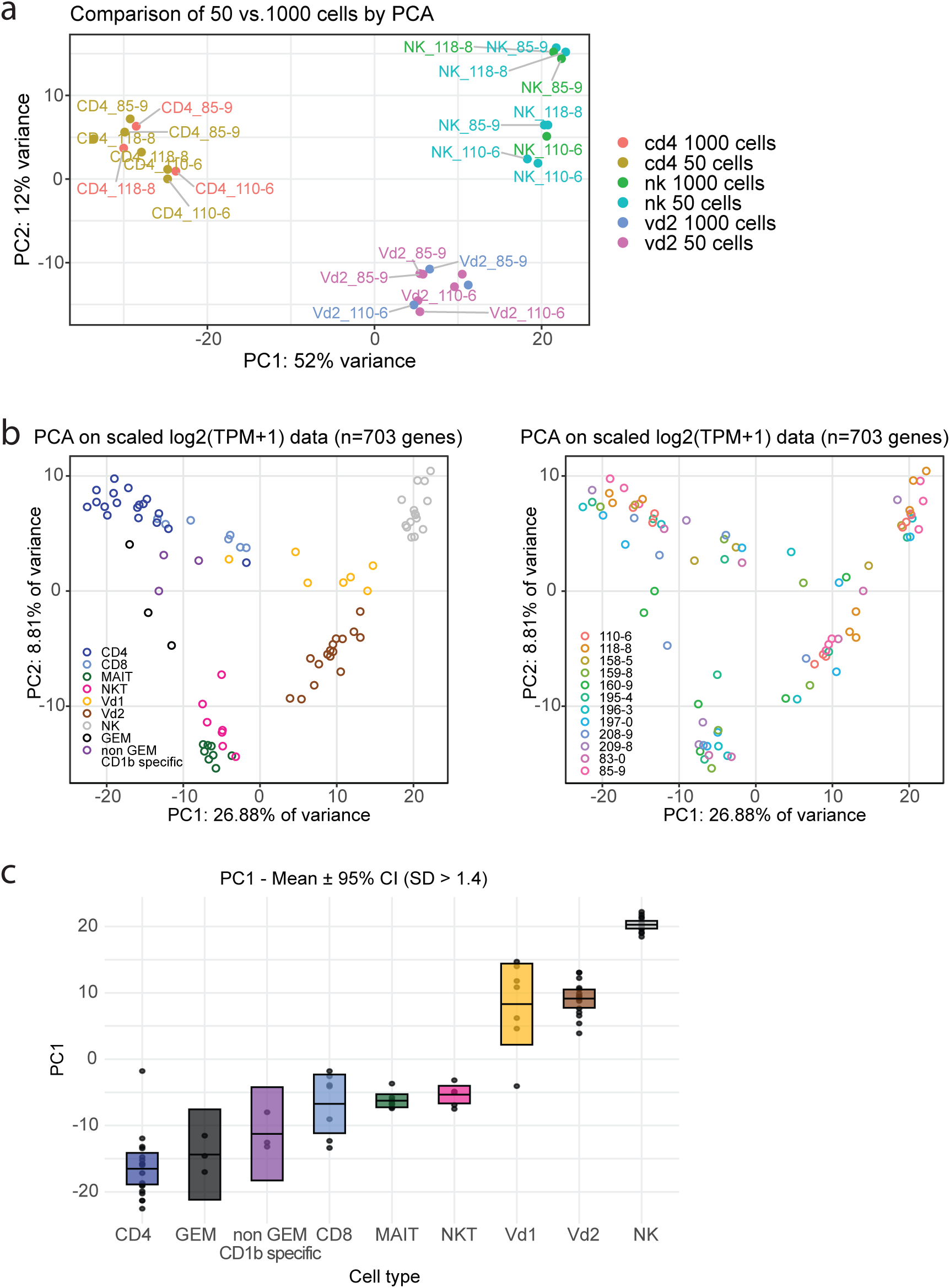
Principal component analysis of transcriptomes. **A-C** PCA was performed on all significantly variable genes. Plotted are scores for PC1 and PC2. **A** PCA plot of technical control samples of 50 or 1000 cells of the same population, and replicate samples of 50 cells. **B** PCA plot of experimental samples colored by cell type (left) or subject (right). **C** PC1 scores by cell type.

### CD1b-specific cells score low on the innateness gradient

After validating the quality of tetramers and sorts, we compared the transcriptional profiles of the two CD1b tetramer^+^ GEM and non-GEM T cell subsets and known innate-like (MAIT, iNKT, Vδ2^+^, Vδ2^−^), adaptive (tet^−^CD4^+^ and CD8^+^) T cell subsets, and NK cells. We excluded the 18 samples consisting of 50 cells from the three participants that were used for the reproducibility and effect of population size study (**Figure 3A)**, and the samples that did not pass quality control due to low read numbers. Of the 96 sequenced samples, 12 were excluded due to low sequencing depth (<500,000 read pairs, compared with >2.6 million for retained samples), consistent with poor RNA quality. The excluded samples comprised all subsets from donor 157-5 and the CD4, CD8, and NK subsets from donor 160-9. To assess the global distribution of samples based on their transcriptomic profiles, we performed principal component analysis (PCA) using the same feature selection strategy as the original analysis of the innateness gradient by filtering for genes with log₂ (transcripts per million (TPM) + 1) > 1 and a standard deviation > 1.4 across samples, yielding 703 genes (**Figure 3B)**. The genes contributing to the loadings of the first and second PCs are listed in **Supplementary Figure 10**. Principal component 1 (PC1) separated the lymphocyte subsets similarly to the previously described “innateness gradient” with conventional CD4^+^ and CD8^+^ T cells on one side (the adaptive side), γδ T cells, MAIT cells, and NKT cells occupying intermediate positions, and NK cells on the innate end (6). GEM T cells had a PC1 score that is close to conventional CD4 T cells, and non-GEM CD1b-specific T cells have a PC1 score in between conventional CD4 and CD8 T cells (**Figure 3B)**. Samples from different donors were intermixed within each cell type, indicating that transcriptional differences between cell types contributed more strongly to PC1 and PC2 separation than donor-specific variation. A quantification of PC1 score per cell type is shown in **Figure 3C**. Thus, relative to the previously reported and currently observed positions of MAIT and NKT cells along the innateness gradient, GEM- and non-GEM CD1b-specific T cells occupy a less innate position along the innateness gradient. However, although GEM T cells and non-GEM CD1b-specific T cells are not separated from conventional T cells across PC1, PC2 moderately separates them.

To gain insight into the specific functions of CD1b-specific T cells, we first examined differences between conventional CD4 T cells and GEM T cells, since the two subsets clustered closely on the PCA plot (**Figure 4A and Supplementary Table 1)**. The differentially expressed gene (DEG) with the highest fold change and the highest significance was TRAV1-2, which was expected because GEM T cells are defined by and were sorted based on TRAV1-2 expression, in addition to CD4 coreceptor expression, and CD1b tetramer staining (**Figure 1)**. The high gene expression of TRAV1-2 provides independent confirmation of the effectiveness of the cell sorting strategy. TRBV6-2 was the only TCR β chain gene in the list, which was reported previously to be enriched among GEM T cells (17, 25), providing two TCR based validations that the cell type of interest was the source of our biologically relevant transcripts. Among genes that are up in GEM T cells, which could potentially positively characterize GEM T cells, there were remarkably few non-TCR genes with a known T cell-specific function: RARG (*p-adj=*0.0016), RORC (*p-adj=*0.013), and TNFRSF18 (GITR) (*p-adj=* 0.024). RARG is one of the retinoic acid nuclear receptors, which has an opposite effect to RORC on Th17 differentiation, and can work in concert with other retinoic acid receptors like RARA in gut homing programs. However, these other retinoic acid receptors are not among the DEGs. Overall, these results do not suggest that GEM T cells form a functionally uniform subset that is clearly distinct from conventional CD4 T cells.

**Figure 4:**
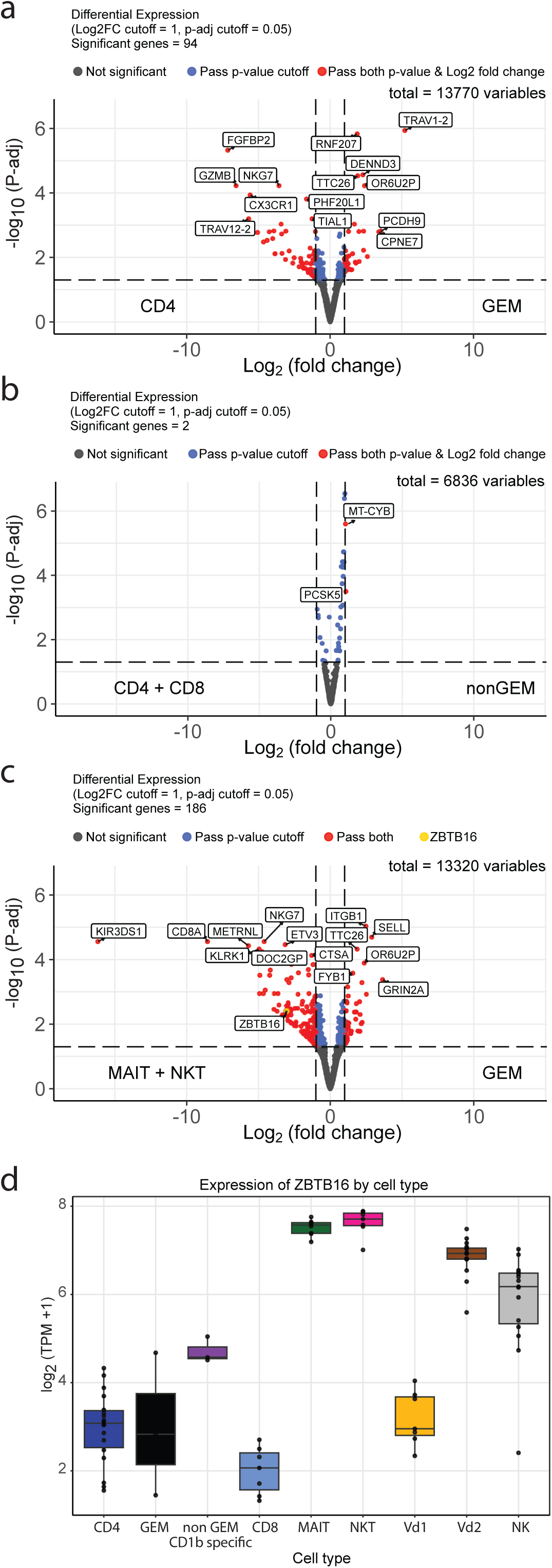
Differential expression of genes in CD1b-specific- and conventional T cells. Volcano plots depicting differentially expressed genes in GEM T cells and conventional CD4 T cells (**A**), non-GEM CD1b-specific T cells and conventional CD8 T + CD4 T cells (**B**), or GEM T cells and the previously studied innate-like T cell populations NKT and MAIT cells (**C**). Dots on the right side represent genes with higher expression levels in GEM T cells or non GEM CD1b-specific T cells.

Since non-GEM CD1b-specific T cells contained both CD4 and CD8 T cells (**Supplementary Figure 11)**, we compared the non-GEM CD1b-specific T cells with all conventional T cells (combining tetramer^−^ CD8 and tetramer^−^ CD4 T cells) (**Figure 4B)**. This comparison was also motivated by the close clustering of CD4 or CD8 and non-GEM CD1b-specific T cells and PC1 score similar to CD4 and CD8 cells. Of note, only 2 DEG were detected at the border of the >2-fold change, without known T cell-specific function. In contrast with the CD4 versus GEM T cell comparison, no TCR variable segment genes uniquely identified non-GEM CD1b-specificspecific T cells. This was expected, as the non-GEM T cell population likely comprises heterogeneous clonotypes and multiple subpopulations with distinct TCR biases. The lack of distinct transcriptional programs for both GEM and non-GEM CD1b-specific T cells again argues against the notion that CD1b-mycolipid-specific cells are a functionally distinct T cell subset.

Lastly, to specifically look for how CD1b-specific T cells differ from established innate-like T cells, we compared (GEM + non-GEM CD1b-specific T cells) with (MAIT + NKT cells) (**Figure 4C**). Of note, we analyzed expression of the Zinc Finger and BTB Domain Containing 16 (ZBTB16), encoding the innate transcription factor Promyelocytic Leukemia Zinc Finger (PLZF), known to be expressed by MAIT and NKT cells (37–40). PLZF expression was significantly in both GEM and non-GEM CD1b-specific T cells (**Figure 4C-D**), further supporting the notion that CD1b-mycobacterial lipid-specific T cells are not closely resembling innate-like T cells. Collectively, the absence of canonical innate-like markers in both the CD4 versus GEM T cell comparison and the comparison of non-GEM CD1b-specific T cells with conventional T cells is consistent with their position along PC1 of the innateness gradient.

## Discussion

In this study, we sought to define how the transcriptional programs of CD1b-specific T cells in the peripheral blood compare to innate-like and adaptive T cell populations. We hypothesized that CD1b-mycolipid-specific GEM T cells, which express an invariant TCR α chain, and possibly also non-GEM CD1b-mycolipid-specific T cells, would functionally resemble other innate-like T cells, including MAIT and NKT cells. Surprisingly, our findings demonstrate that CD1b-specific T cells are transcriptionally distinct from known innate-like T populations and instead resemble conventional adaptive CD4 T cells. In line with this conclusion, both CD1b-specific subsets also lack expression of PLZF, a transcription factor that predisposes innate-like T cells to rapid effector functions. The observed divergence from innate-like T cell populations opens the possibility that GEM T cells and non-GEM CD1b-specific T cells follow a more conventional T cell life cycle. If this is true, we would predict that GEM T cells and non-GEM CD1b-specific T cells are poised to follow more conventional T cell differentiation trajectories from naïve to effector and memory phenotypes.

In a previous study, we did not observe significant differences in the peripheral blood frequencies of GEM- and non-GEM CD1b-specific T cells between persons with TB disease and their asymptomatic household contacts (24). However, both groups exhibited generally higher frequencies than healthy blood bank donors without prior mycobacterial exposure, consistent with their memory potential (24). Our current study did not separate naïve, activated, and memory T cell populations of the GEM T cells and non-GEM CD1b-specific T cell populations due to the limited cell numbers and available PBMCs. Nonetheless, we previously showed that both GEM and non-GEM CD1b-GMM tetramer^+^ T cells have a higher proportion of CD45RO expressors than their tetramer-negative counterparts, and in vivo clonal expansion can be detected in some people (17, 24). Clonal expansion of GEM T cells was also reported by others in a South African cohort (25). Together, these findings suggest that GEM T cells and non-GEM CD1b-specific T cell populations follow a more conventional life cycle. Should this be true, it would also be expected that GEM and non-GEM CD1b-specific T cells are diverse in their final differentiation outcomes, just like conventional CD4 T cells, which is consistent with the absence of a single clear Th or other functional profile in our data. The following limitations should be considered when interpreting our findings. First, our study was limited by a small sample size and low frequency of detectable CD1b-specific GEM and non-GEM T cells across donors, which constrained statistical power and potentially under-reported appreciable transcriptional programs that uniquely define these cells. In addition, selecting individuals with sufficient cell numbers for sorting may have inherently enriched for expanded clones with distinct transcriptional programs. Finally, we did not include a mycobacteria-naïve control group, limiting our ability to determine the extent to which the observed transcriptional features are shaped by prior mycobacterial exposure. Regardless, our findings are consistent with the conceptual framework proposed by Lepore et al. (41), who coined the term “adaptive-like” to describe unconventional T cells that recognize non-peptide antigens presented by non-polymorphic molecules, yet share functional characteristics with conventional adaptive MHC-specific T cells.

We do not, however, propose that our results are generalizable to all CD1a, CD1b, and CD1c-specific T cells. One consideration is that the populations that were studied here, GEM and non-GEM CD1b-specific T cells, co-recognize CD1b and a rare mycobacterial lipid antigen that is not usually encountered in the human body. Thus, CD1b-specific T cells may conform to the “needle in a haystack” paradigm that was originally proposed for MHC-peptide-specific T cells, with a low naïve precursor frequency, which expands upon encountering the cognate lipid antigens. This is in sharp contrast with the many autoreactive CD1-specific T cells that have been isolated from human blood (21, 42–44) and tissues, like the skin (45–47), for which innate-like functions are still being considered. In this setting, it has been proposed that expression of the relevant CD1 isoforms acts as a danger signal or a homeostatic signal. This latter category of CD1-specific T cells is likely to be transcriptionally closer to the innate side of the innateness gradient than GEM and non-GEM CD1b-specific T cells.

One aspect of the field that remains difficult to reconcile with our finding is the known difference in thymic selection between MHC-specific T cells and CD1-specific T cells and their proposed implication for T cell development. For murine NKT cells, it is known that they are positively selected on (cortical) CD4^+^CD8^+^ double-positive thymocytes instead of cortical thymic epithelial cells. It is thought that this distinct selection mechanism drives the expression of PLZF due to homotypic interactions between Signaling Lymphocytic Activation Molecule (SLAM) proteins on the thymocyte that receives a concomitant TCR signal and SLAM on the thymocyte that expresses CD1d or MR1 for iNKT or MAIT cells, respectively. PLZF is considered the master switch for innate-like effector functions (37–40, 48). Human thymocytes express very high levels of all CD1 isoforms, so it is thought that all CD1a, CD1b, CD1c, and CD1d-specfic cells are selected by thymocytes. Because all thymocytes express SLAM, this would be predicted to bias all CD1-specific T cells towards expression of PLZF and innate gene sets. The absence of PLZF and innate profiles in GEM T cells and non-GEM CD1b-specific T cells in our data is at odds with this prediction, which suggest that maybe not all SLAM-SLAM interactions lead to PLZF expression, or early PLZF expression might be lost and not necessarily lead to innate-like T cell profiles. Related, unsolved questions in the field concern the lipid antigens that select CD1-specific T cells in the thymus. This question is partly answered for NKT cells, but not for mycobacterial lipid-specific populations like GEM T cells and non-GEM CD1b-specific T cells that we studied here. Overall, the selection of CD1a, CD1b, and CD1c-specific T cells in the thymus is an understudied area, which is mostly due to the absence of CD1a, CD1b, and CD1c in wild-type mice, the relatively low precursor frequency of antigen-specific CD1a-, CD1b-, and CD1c-specific T cells, and the difficulties with obtaining and performing mechanistic studies in human thymus biospecimens.

## Author contributions (with CRediT details)

Alan Hsieh: Formal analysis, Investigation, Visualization; Kattya Lopez: Formal analysis, Investigation, Visualization; Segundo R León: Resources; Roger I Calderon: Resources; Leonid Lecca: Resources; Megan B. Murray: Funding acquisition, Resources; D. Branch Moody: Conceptualization, Funding acquisition, Methodology, Supervision, Writing – review & editing; Sara Suliman: Conceptualization, Data curation, Formal analysis, Methodology, Supervision, Visualization, Writing – original draft, Writing – review & editing; Ildiko Van Rhijn: Conceptualization, Data curation, Formal analysis, Methodology, Supervision, Visualization, Writing – original draft, Writing – review & editing.

Choose from: Conceptualization, Data curation, Formal analysis, Funding acquisition, Investigation, Methodology, Project administration, Resources, Software, Supervision, Validation, Visualization, Writing – original draft, Writing – review C editing

## Data availability

All raw RNA-seq data are deposited into Gene Expression Omnibus (GEO) Repository, Accession number: XXXXX.

