## Supplemental Figures for "CD1b-specific T cells are transcriptionally closer to conventional CD4 T cells than to innate-like NKT and MAIT cells"

### Figure S1

subject 083-0

first sort on PBMC sample:

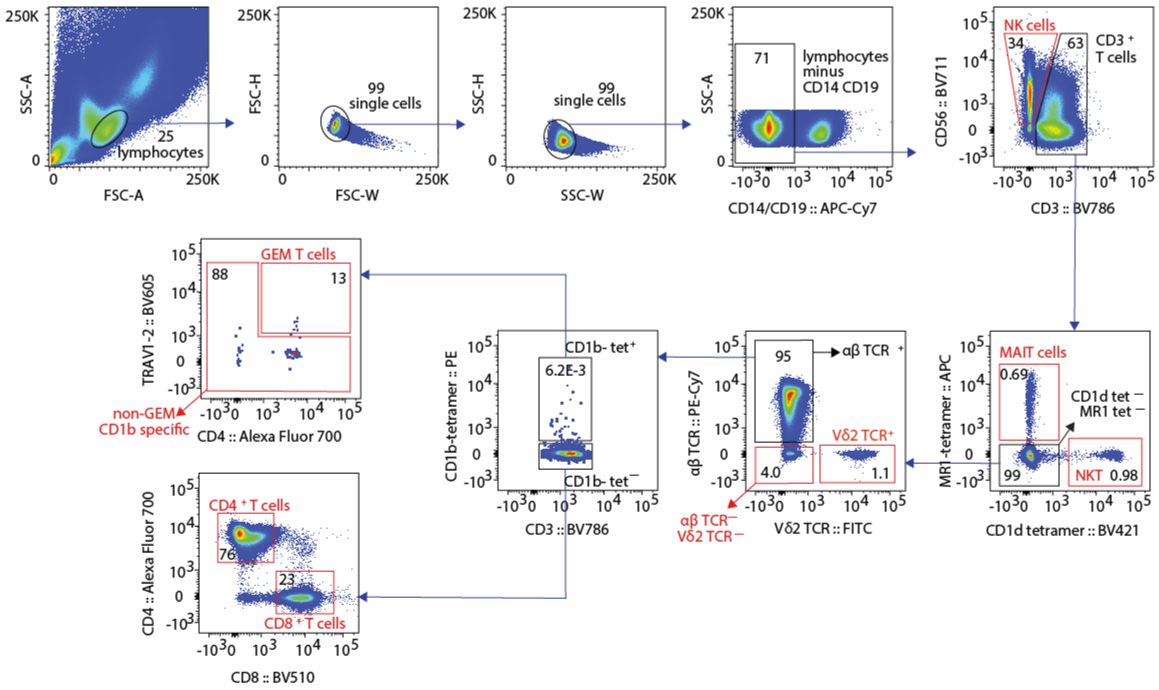


pre second sort: cell populations

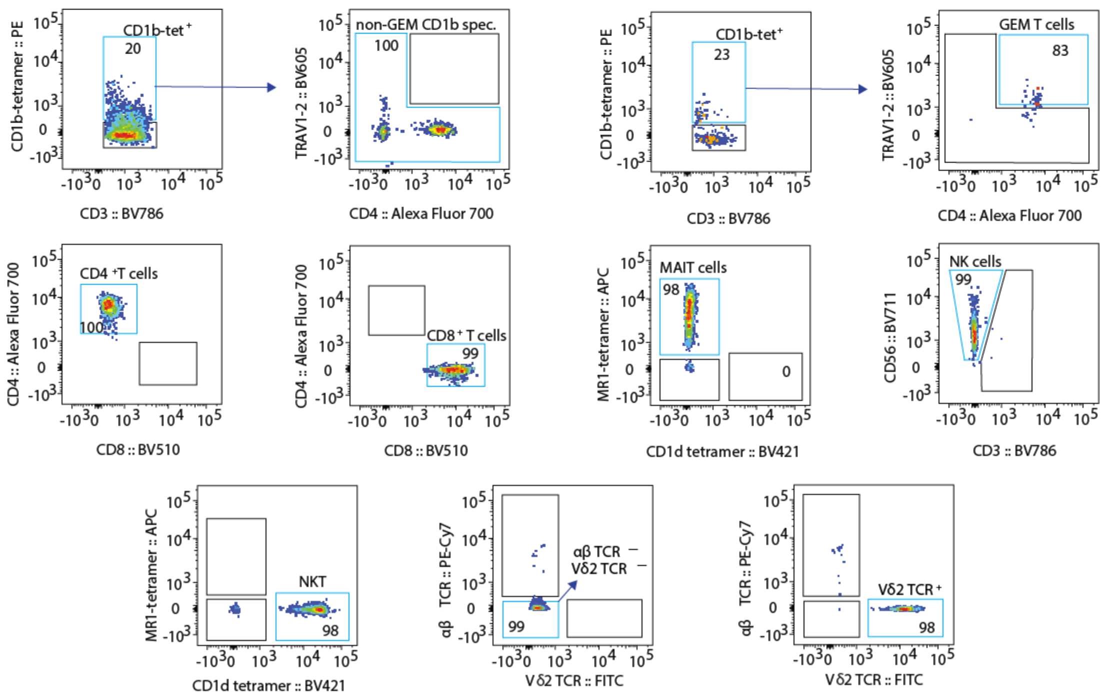

### Figure S2

subject 157-5

first sort on PBMC sample:

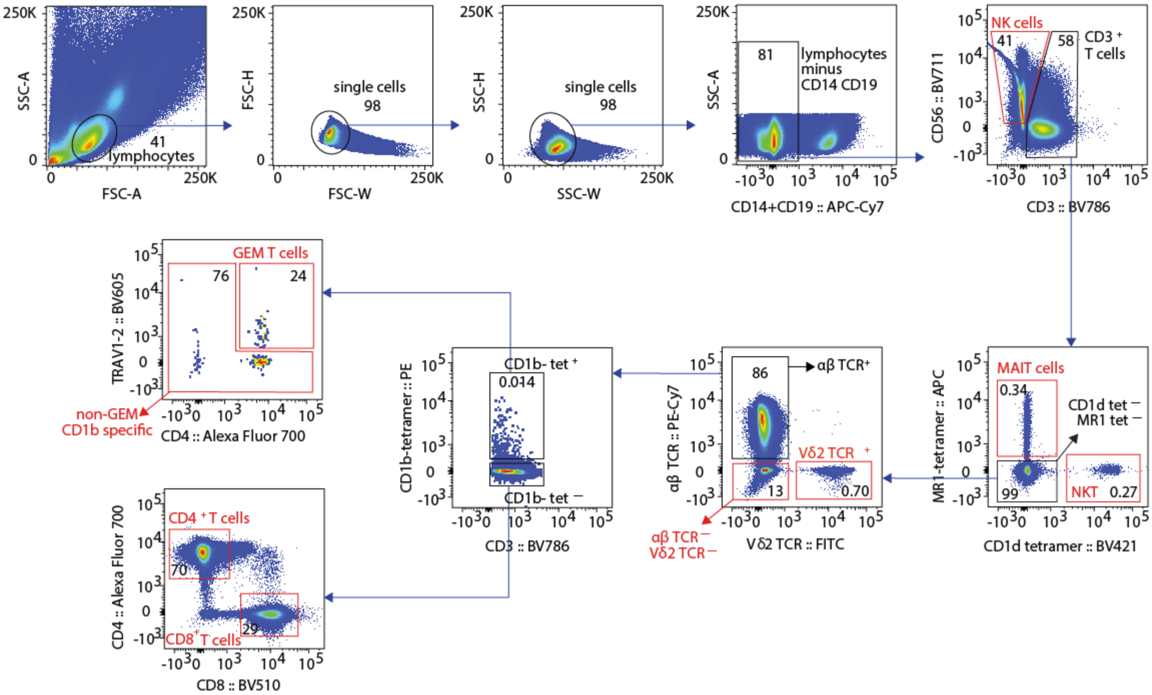


pre second sort: cell populations

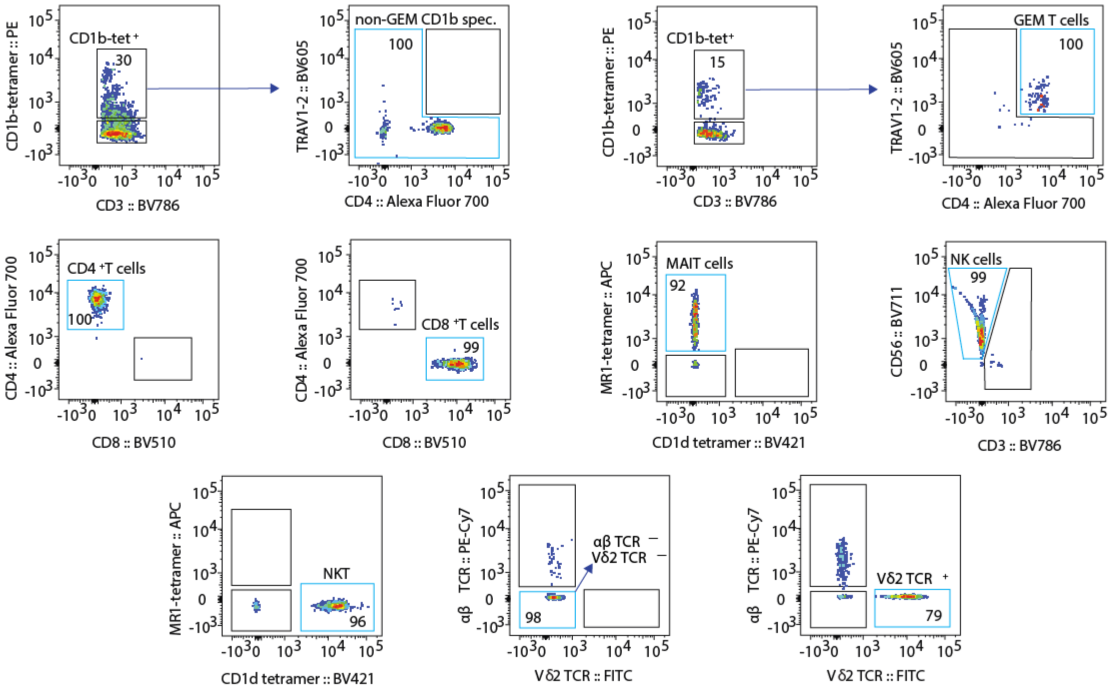

### Figure S3

donor 158-5

first sort on PBMC sample:

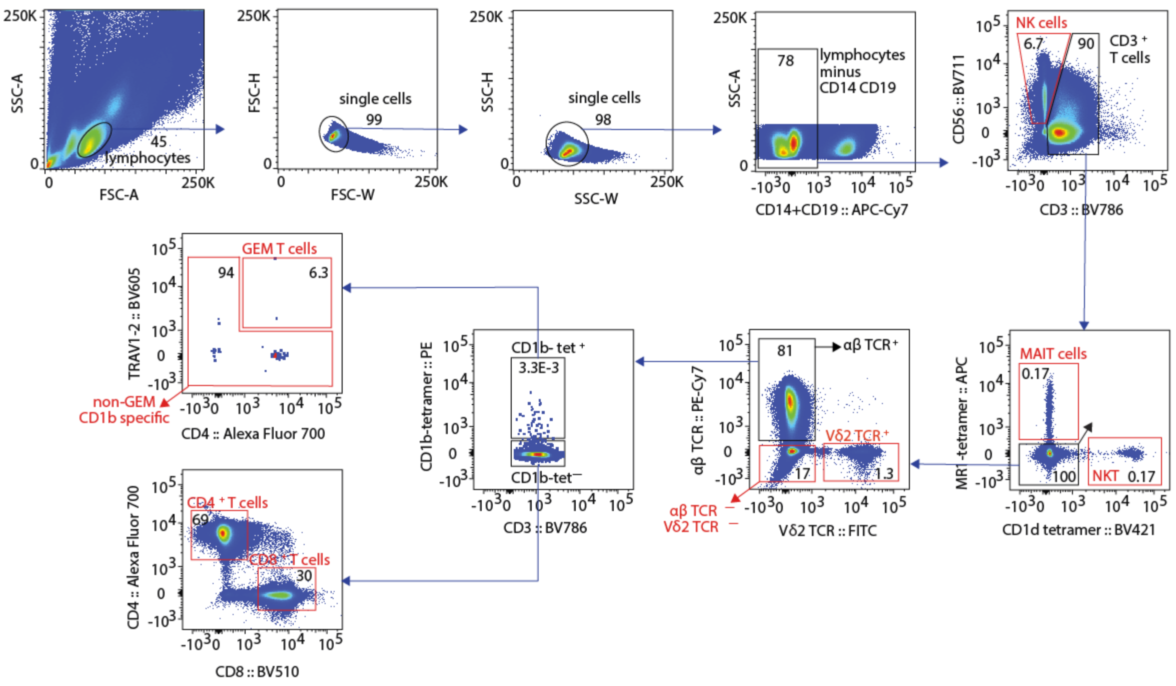


after first sort: cell populations

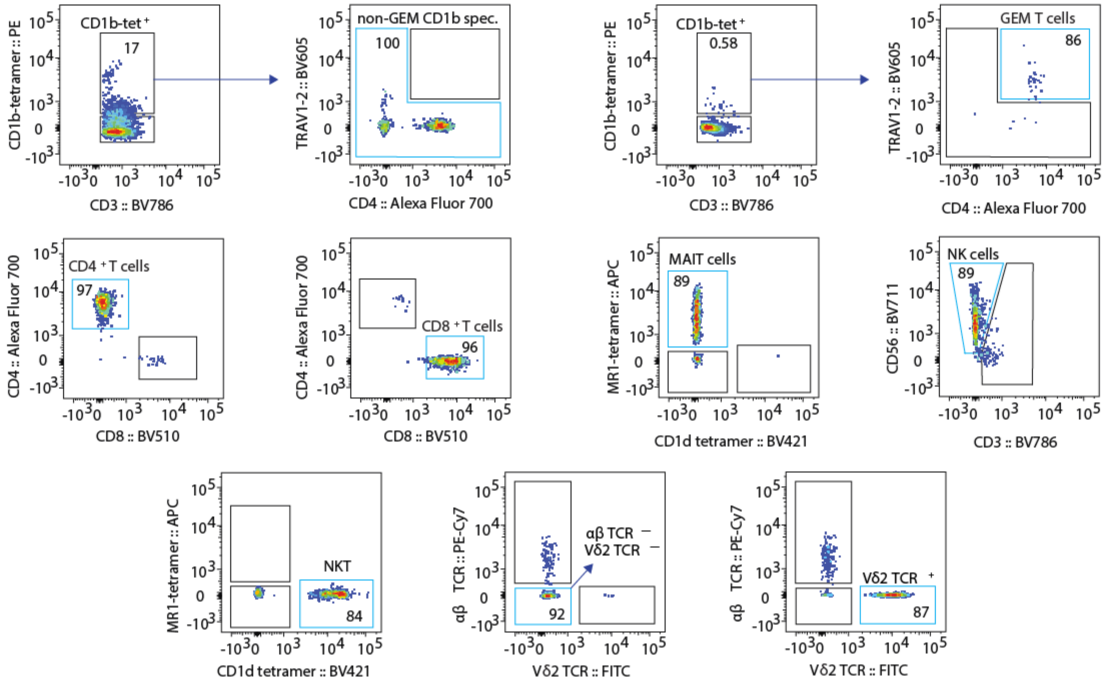

### Figure S4

subject 159-8

first sort on PBMC sample:

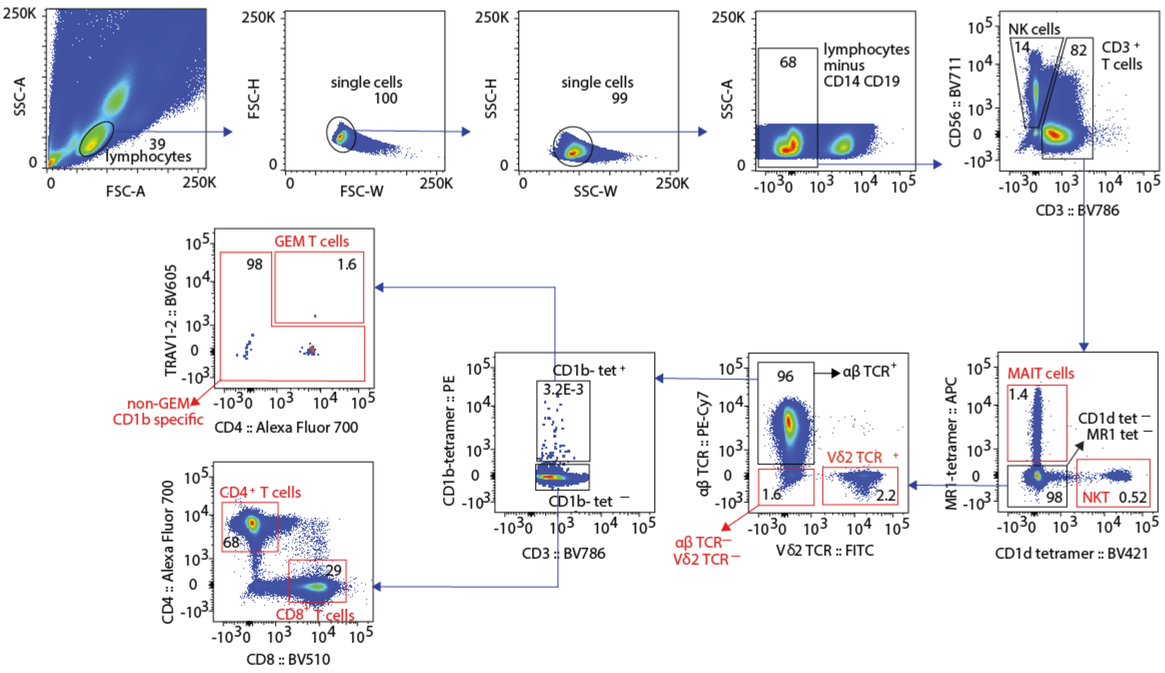


pre second sort: cell populations

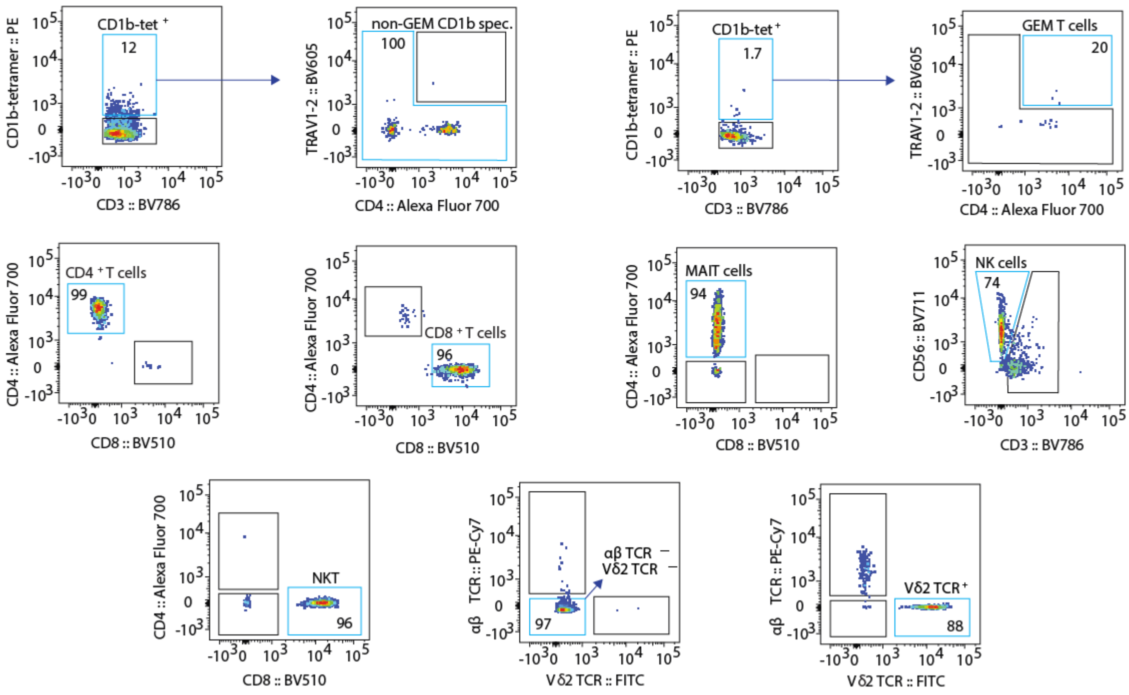

### Figure S5

subject 160-5

first sort on PBMC sample:

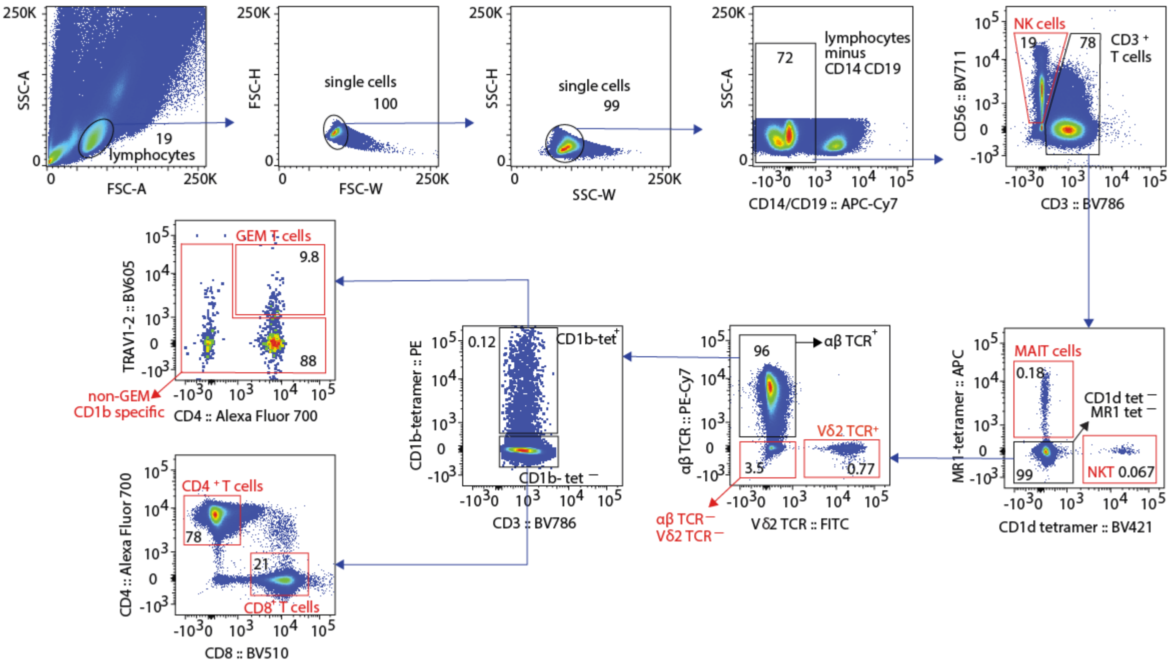


pre second sort: cell populations

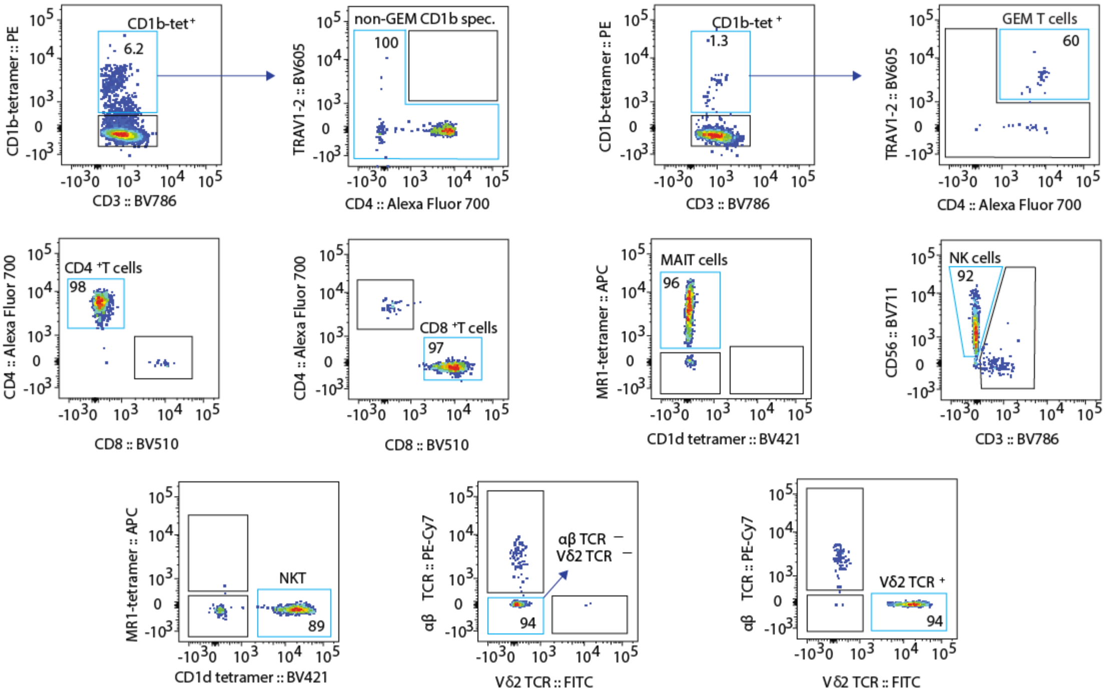

### Figure S6

subject 195-4

first sort on PBMC sample:

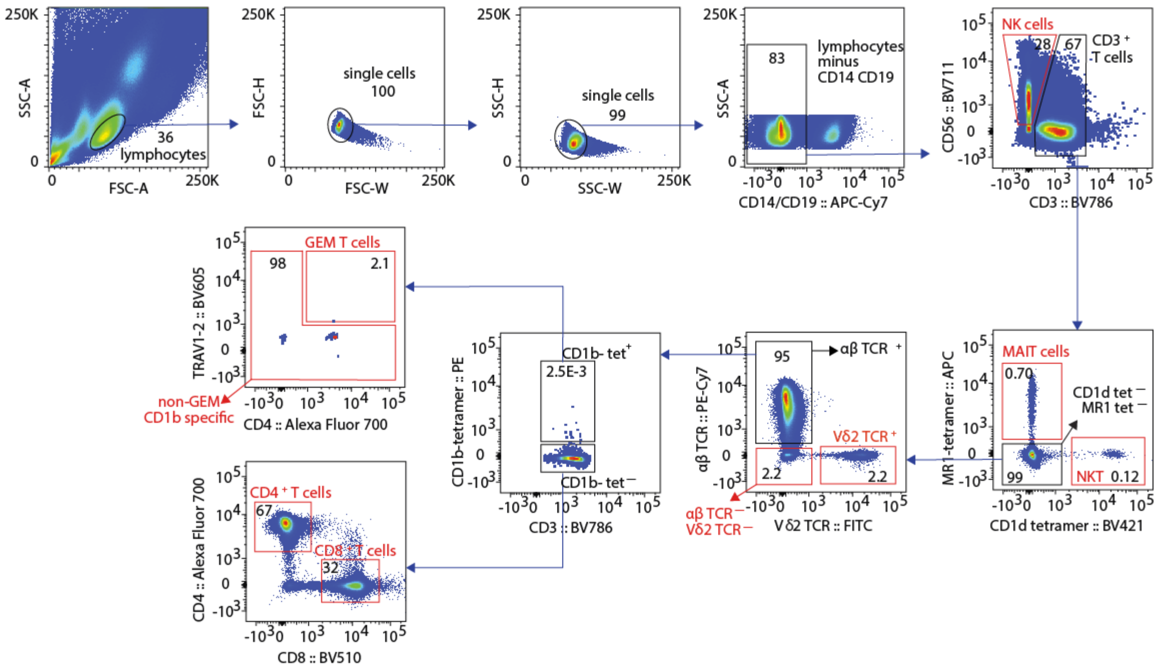


pre second sort: cell populations

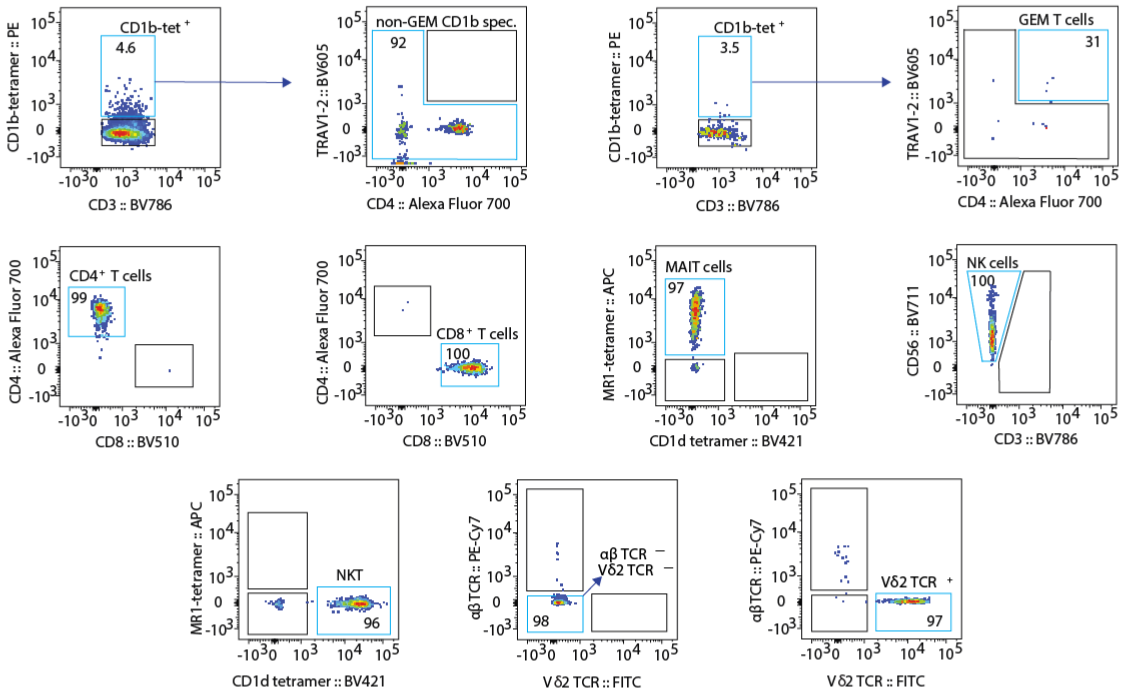

### Figure S7

subject 196-3

first sort on PBMC sample:

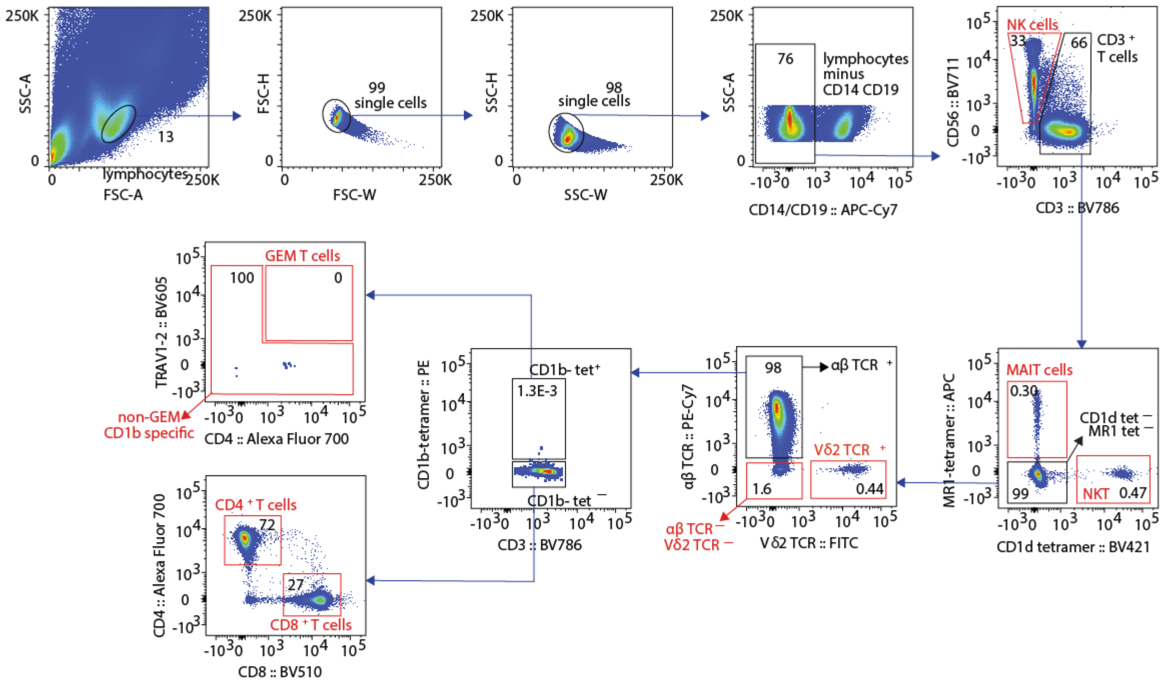


pre second sort: cell populations

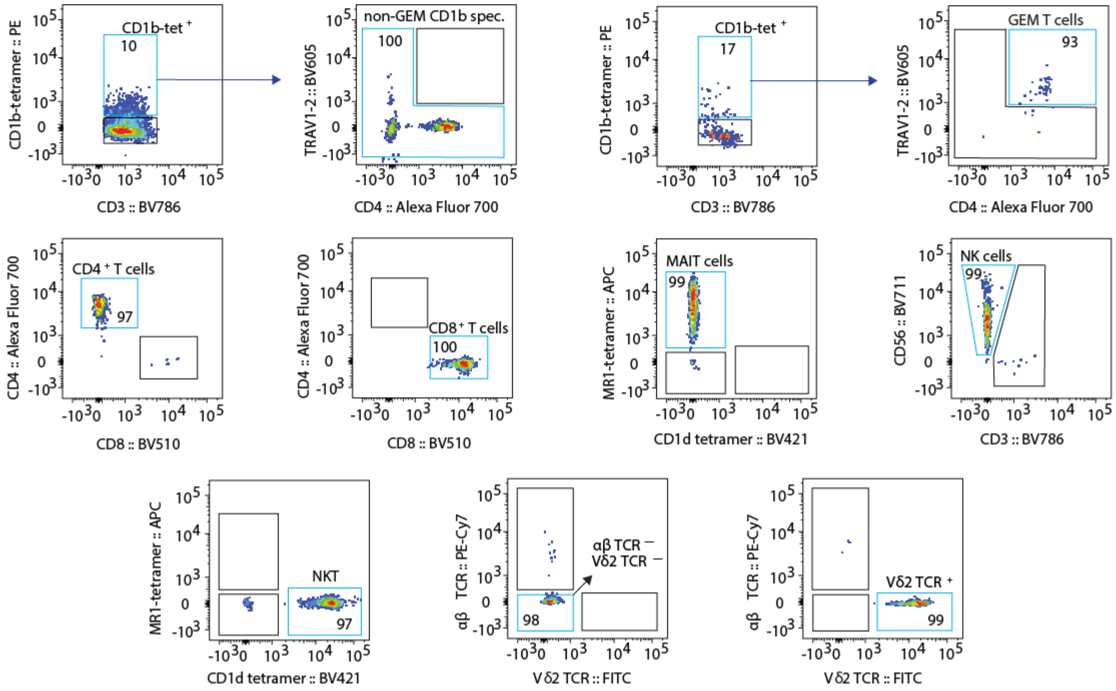

### Figure S8

subject 197-0

first sort on PBMC sample:

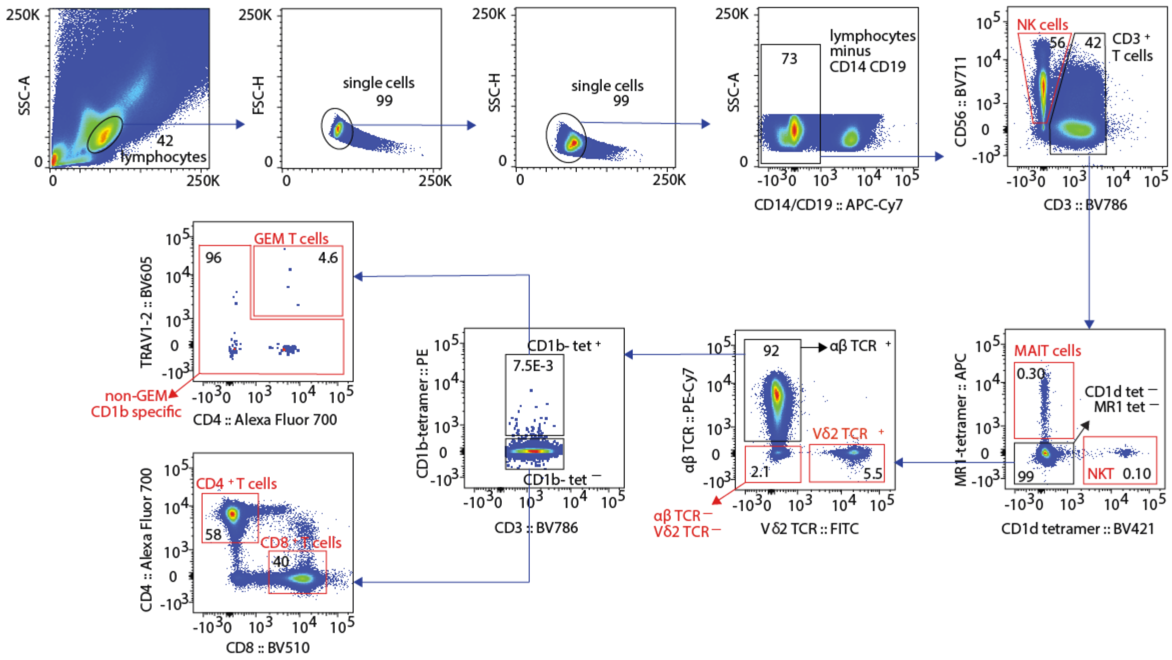


pre second sort: cell populations

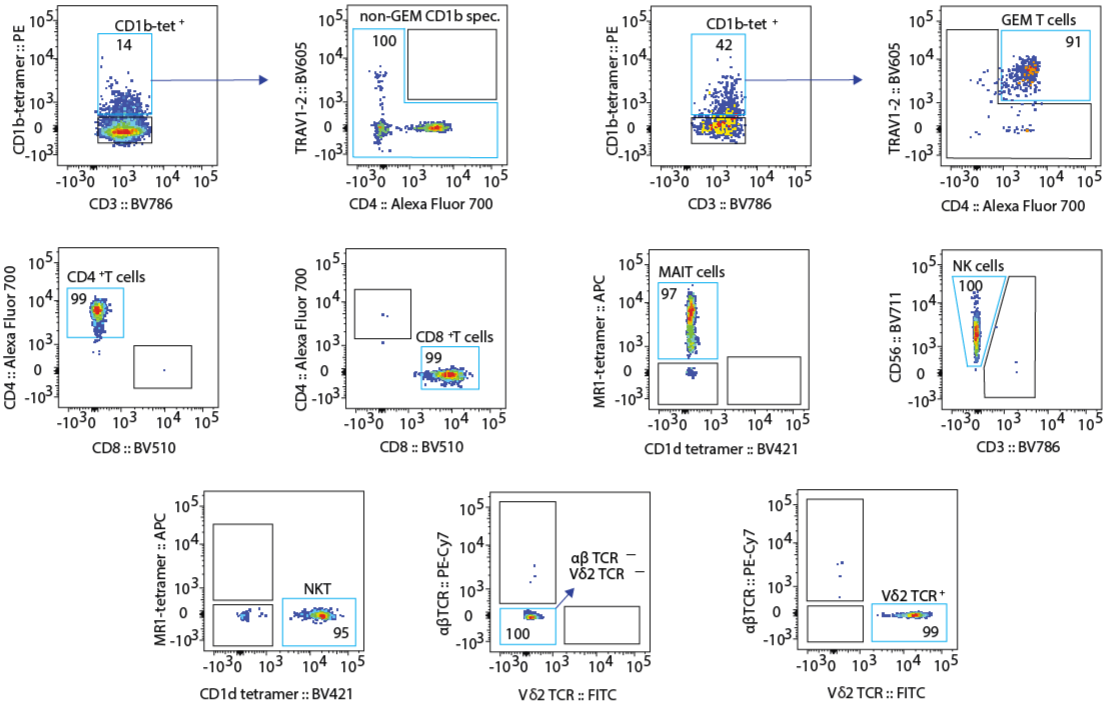

### Figure S9

subject 209-8

first sort on PBMC sample:

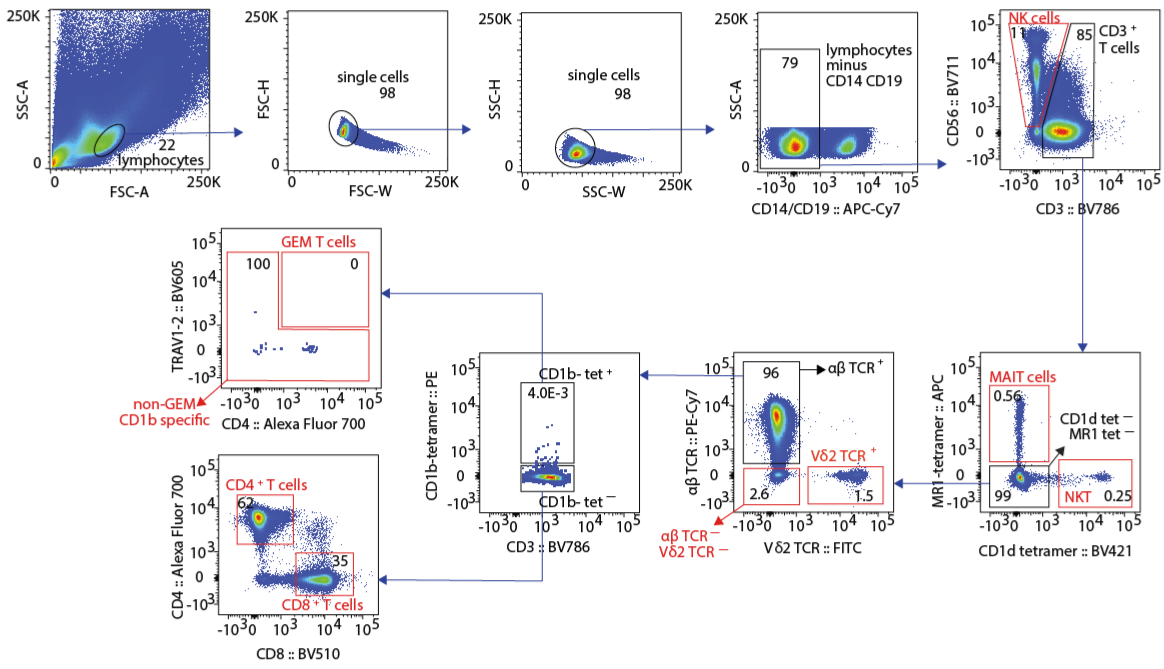


pre second sort: cell populations

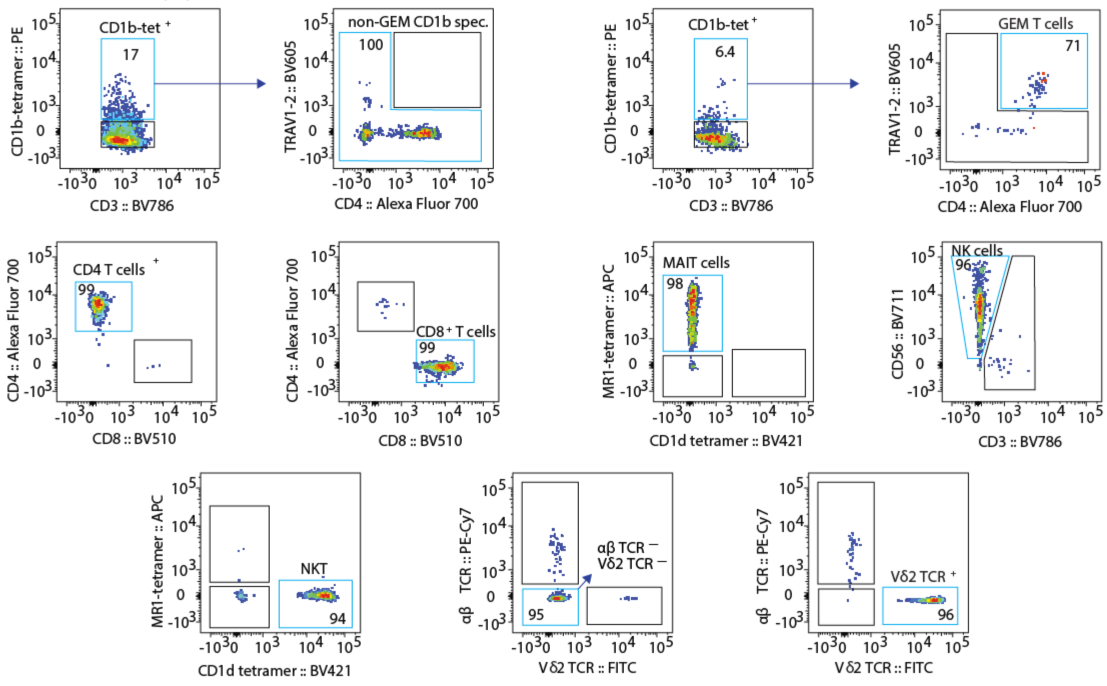

Figure S10

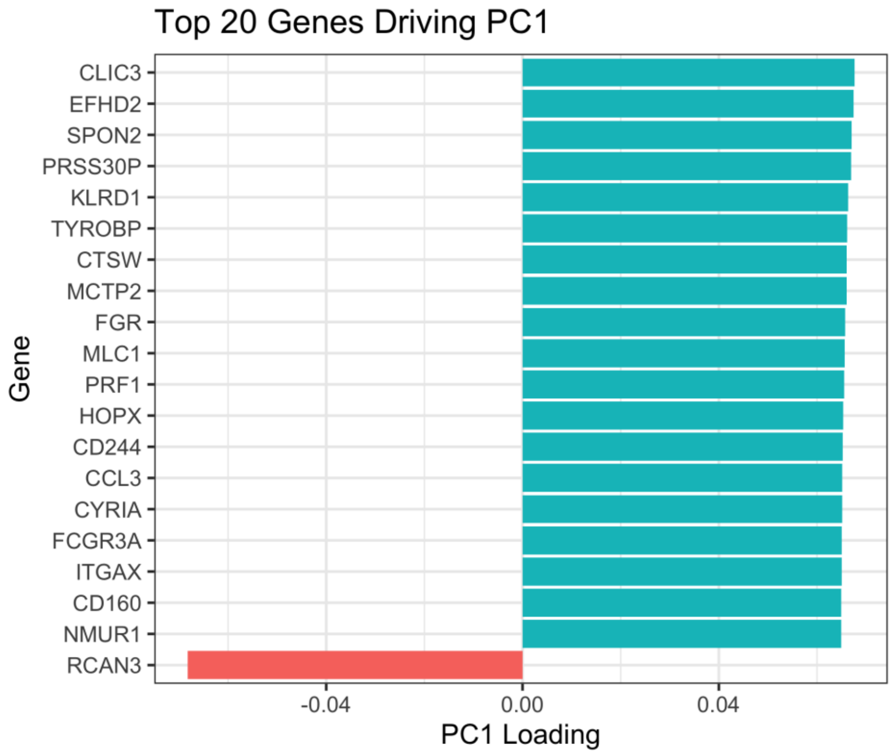

Top 20 Genes Driving PC1
Gene
CLIC3
EFHD2
SPON2
PRSS30P
KLRD1
TYROBP
CTSW
MCTP2
FGR
MLC1
PRF1
HOPX
CD244
CCL3
CYRIA
FCGR3A
ITGAX
CD160
NMUR1
RCAN3
-0.04
0.00
0.04
PC1 Loading


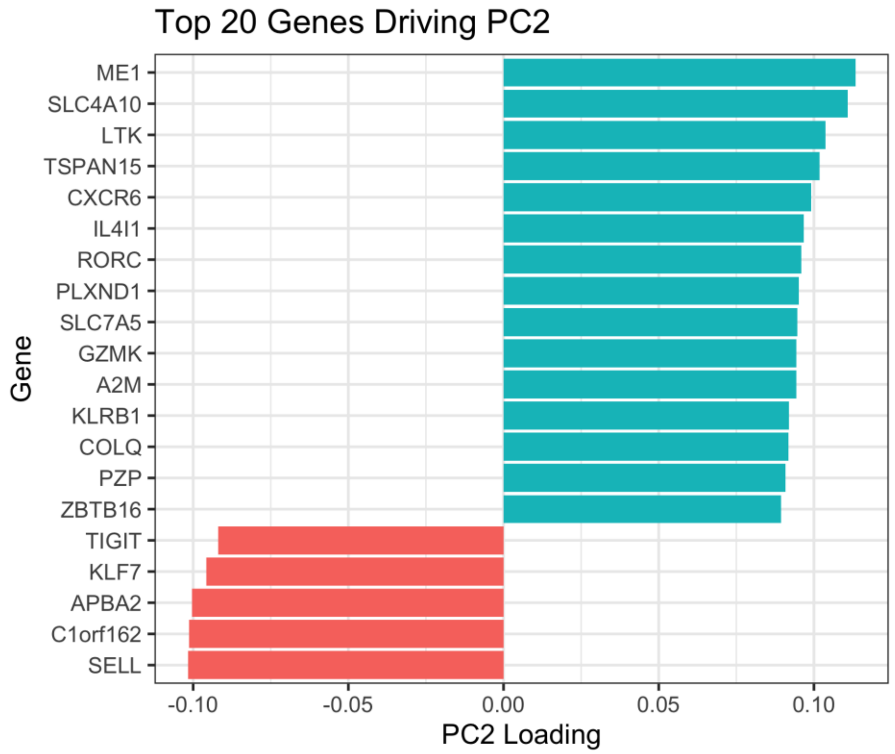

Top 20 Genes Driving PC2
Gene
ME1
SLC4A10
LTK
TSPAN15
CXCR6
IL4I1
RORC
PLXND1
SLC7A5
GZMK
A2M
KLRB1
COLQ
PZP
ZBTB16
TIGIT
KLF7
APBA2
C1orf162
SELL
-0.10
-0.05
0.00
0.05
0.10
PC2 Loading

### Figure S11

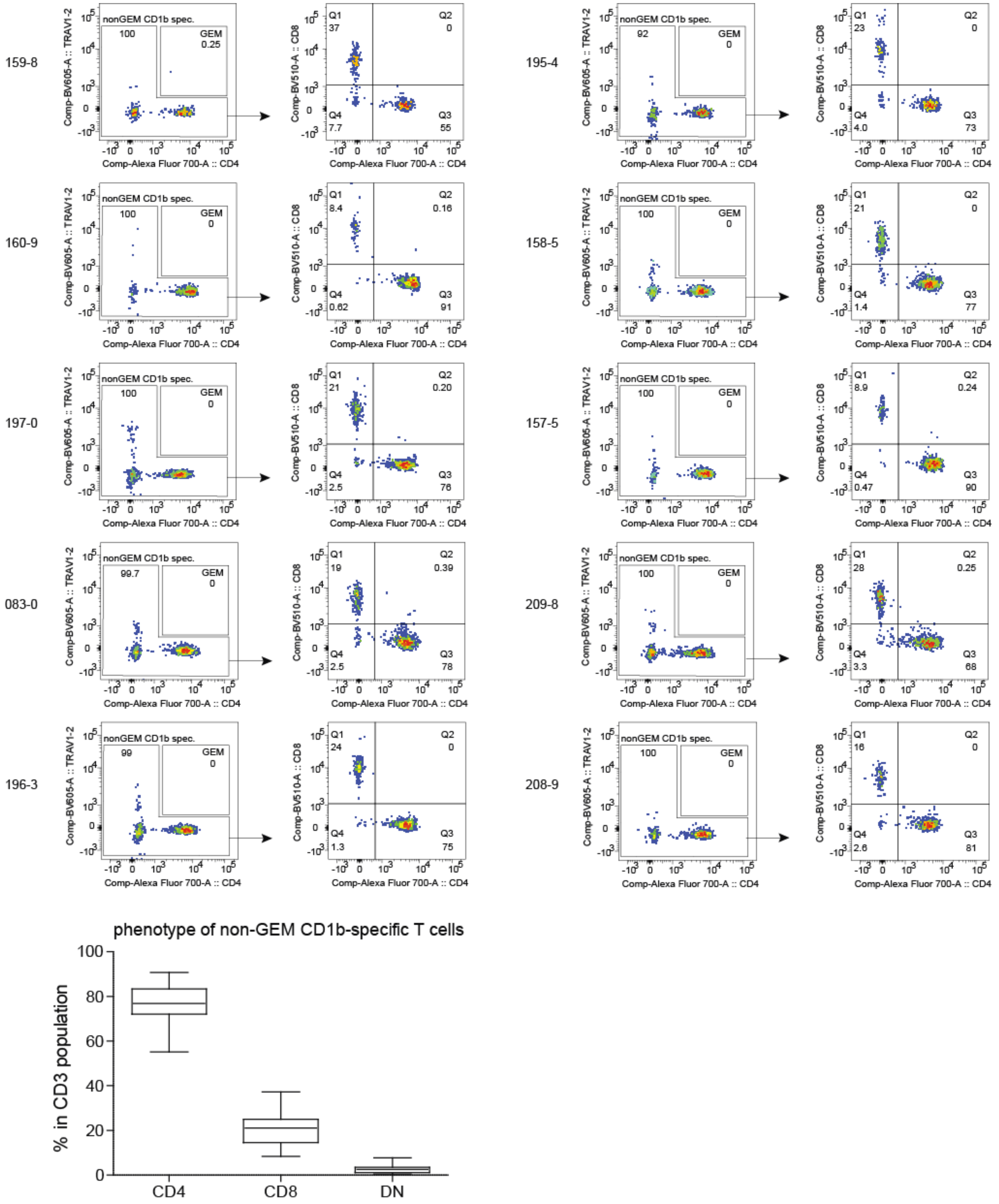

159-8
nonGEM CD1b spec.
100
GEM
0.25
Q1 37
Q2 0
Q4 7.7
Q3 55
195-4
nonGEM CD1b spec.
92
GEM
0
Q1 23
Q2 0
Q4 4.0
Q3 73
160-9
nonGEM CD1b spec.
100
GEM
0
Q1 8.4
Q2 0.16
Q4 0.62
Q3 91
158-5
nonGEM CD1b spec.
100
GEM
0
Q1 21
Q2 0
Q4 1.4
Q3 77
197-0
nonGEM CD1b spec.
100
GEM
0
Q1 21
Q2 0.20
Q4 2.5
Q3 76
157-5
nonGEM CD1b spec.
100
GEM
0
Q1 8.9
Q2 0.24
Q4 0.47
Q3 90
083-0
nonGEM CD1b spec.
99.7
GEM
0
Q1 19
Q2 0.39
Q4 2.5
Q3 78
209-8
nonGEM CD1b spec.
100
GEM
0
Q1 28
Q2 0.25
Q4 3.3
Q3 68
196-3
nonGEM CD1b spec.
99
GEM
0
Q1 24
Q2 0
Q4 1.3
Q3 75
208-9
nonGEM CD1b spec.
100
GEM
0
Q1 16
Q2 0
Q4 2.6
Q3 81
Comp-BV605-A :: TRAV1-2
Comp-BV510-A :: CD8
Comp-Alexa Fluor 700-A :: CD4
phenotype of non-GEM CD1b-specific T cells
% in CD3 population
100
80
60
40
20
0
CD4
CD8
DN
